# Perceptual integration of multisensory haptic, visual, and auditory feedback for roughness discrimination in augmented reality

**DOI:** 10.64898/2026.09.09.748749

**Authors:** Sarah Bonnet, Rebecca Ericsson, Iuliia Zhurakovskaia, Heidy Daumas, Rochelle Ackerley

## Abstract

Understanding how our different senses interact to shape our perception is essential to design realistic and immersive virtual and augmented reality (VR/AR) experiences. The present study investigated how roughness perception can be modulated through haptic, visual, and auditory cues in AR using a vibrotactile wristband. Participants compared virtual textures varying in vibration frequency/amplitude, visual grain size, and friction sound. Results revealed strong linear relationships between stimulus parameters and perceived roughness, with haptic frequency and visual cues driving the highest discrimination performance. Adding non-informative sensory feedback reduced perceptual sensitivity, acting as noise. Individual differences emerged: participants who rated haptic as the easiest modality showed greater sensitivity to haptic variations, while visual-reliant participants performed better with visual cues. We conclude that roughness in AR can be systematically manipulated, but is vulnerable to perceptual interference from irrelevant inputs, where our work provides actionable insights for implementing optimized and adaptive AR/VR interfaces.

## INTRODUCTION

Virtual and augmented reality (VR/AR) technologies are increasingly integrated into diverse fields such as education, healthcare, professional training and gaming (Parekh et al. 2020; Xie et al. 2021). While VR immerses users entirely within a synthetic environment, AR blends virtual elements into the real world, allowing for simultaneous interactions with both virtual and physical object. This blending of realities in AR holds great potential for enhancing user experiences, particularly for replicating natural sensory interactions. Among the various sensory modalities involved, touch, and specifically the perception of surface textures, plays a crucial role in the realism and engagement with immersive environments. Roughness perception, a fundamental aspect of tactile exploration (Tanaka et al. 2014), enables individuals to assess surface properties and make judgments critical for interactions and object recognition (Smeets and Brenner 1999; Wang et al. 2024). In everyday environments, this percept can be modified through multisensory integration, involving not only tactile signals, but also visual and auditory cues. Reproducing such rich multisensory experiences in AR remains a challenge, especially given the absence of direct physical feedback from virtual objects. Understanding how roughness perception can be recreated and modulated through artificial sensory feedback is therefore important for advancing the realism and effectiveness of immersive interfaces.

The perception of roughness arises from the contact between the skin and a surface, where, physical variations in surface height generate vibrations in the skin, that are captured by mechanoreceptive afferents (Fradin et al. 2025). Texture perception can be differently experienced over the skin (Ackerley et al. 2014), depending on the type and density of mechanoreceptors present, but our main means for active texture exploration is via the glabrous skin of the fingers. Fast-adapting type 1 (FA1, Meissner corpuscles) and slowly-adapting type 1 (SA1, Merkel cells) are densely represented in the human fingertip, although all low-threshold mechanoreceptors contribute to the awareness of touch (Vallbo and Johansson 1984). Coarse textures, with spatial periods greater than 200 µm, are primarily perceived through the firing of SA1 afferents (Bensmaia and Hollins 2003). In contrast, fine textures with spatial periods below 200 µm, are perceived through vibrotactile channels, such as via the fast-adapting type 2 (FA2, Pacinian corpuscles) afferents, which respond well to high frequency vibrations (Weber et al. 2013). Recent work has further shown that FA1s encode multiple parameters of touch simultaneously, including force, speed, and spatial period (Lang et al. 2025). Building on the importance of vibrations in roughness, a vibrotactile display modulating frequency and amplitude showed that specific patterns systematically change roughness perception (Asano et al. 2014, 2012). Further work investigated how different vibrotactile signals, through variations in frequency, amplitude, and temporal patterns can generate distinct haptic textures, revealing how these parameters shape the perception of roughness, bumpiness, adhesiveness, and sharpness (Strohmeier and Hornbæk 2017).

Recreating texture artificially in AR is challenging due to the absence of direct physical feedback. Wearable devices, such as haptic rings and wristbands, can simulate localized sensations (Gaudeni et al. 2019; Friesen and Vardar 2024), but replicating the full spectrum of real-world textures remains difficult, especially given individual differences in haptic sensitivity and roughness perception (Bergmann Tiest and Kappers 2007). Beyond touch, visual and auditory cues also shape roughness perception. Visual texture properties (e.g. grain size, density; Klatzky & Lederman, 2010) provide predictive information before contact (Klatzky et al., 1993; Whitaker et al., 2008), and visual expectations can even activate tactile central networks in the absence of touch (Sun et al. 2016). These visual expectations can actively modulate tactile perception: prior exposure to visual stimuli has been shown to significantly alter subsequent haptic perception, depending on the material observed beforehand (Yanagisawa and Takatsuji 2015). Similarly, task-irrelevant visual motion cues are sufficient to modify roughness perception, with incongruent visual motion tending to make surfaces feel smoother (Suzuishi et al. 2020). Auditory cues also play a meaningful role in roughness perception. Friction-induced sounds arising from skin-texture interactions depend on factors such as applied pressure, movement speed, and surface spatial properties, and have been shown to modulate or even substitute tactile roughness perception (Lederman 1979; Guest et al. 2002). The classic parchment-skin illusion demonstrates how auditory feedback can alter perceived roughness through changes in sound loudness and frequency (Jousmäki and Hari 1998). Auditory roughness can also emerge as a purely acoustic phenomenon arising from amplitude fluctuations (Vassilakis and Kendall 2010), which can even be elicited vibrotactually, suggesting shared cross-modal perceptual mechanisms (Bernard et al. 2022). Overall, however, auditory inputs tend to be weighted less than haptic or visual information in multisensory roughness integration (Guest and Spence 2003; Klatzky and Lederman 2010a), with vision supporting faster and more accurate categorization, and haptics providing precise information about physical properties.

Studies on virtual texture perception illustrate how visual information can enhance haptic experiences, creating a more immersive experience (Friesen and Vardar 2024). In AR, virtual surfaces inherently lack the mechanical properties of real materials, making artificial reconstruction necessary. While vibrotactile parameters, visual grain size, texture density, and auditory cues can each individually influence texture perception (Guest et al. 2002; Bernard et al. 2022; Ho et al. 2006), few studies have systematically combined these modalities to explore their integrated effects on roughness perception. Moreover, incongruent or task-irrelevant cues may change the sensory experience, as the reliability and congruency of information determine how cues are weighted in multisensory integration (Ernst and Bülthoff 2004). A key methodological consideration in this context concerns the placement of haptic feedback devices. Fingertip-based actuators, while providing direct mechanical stimulation at the primary site of texture exploration, constrain finger movement and prevent natural contact interactions (Friesen and Vardar 2024; Normand et al. 2025). To preserve the full range of finger motion and avoid restricting interactions, a wristband-based vibrotactile approach was adopted in the present study, transmitting vibrations near the hand, while leaving the finger free. Whereas previous work has examined vibrotactile roughness perception with direct fingertip contact on physical surfaces across immersive environments (Normand et al. 2024), the present study investigated roughness perception in AR in the absence of physical contact, a frequent situation where users interact with purely virtual objects, and where inducing convincing tactile sensations without any contact remains a challenge.

The present study aimed to manipulate roughness parameters systematically across haptic, visual, and auditory modalities, via a vibrotactile wristband, through simulating virtual sandpaper textures of varying granularity, and by audio recordings of sandpaper interactions. We formulated three hypotheses: (1) each modality will independently and systematically influence perceived roughness; (2) additional sensory input may act as perceptual noise when it lacks task-relevant information; and (3) individual sensory preferences will modulate participants’ sensitivity to roughness variations across modalities.

## METHODS

### Experimental model and study participant details

For this experiment, 30 healthy human participants were recruited: 15 females, 7 left-handed (self-reported), aged between 19 and 33 years (mean age 25 years). Age and sex were included as covariates in linear regression analyses to assess their potential influence on the results, but no significant effects or associations were identified. All participants read and signed a written informed consent form before participating in the study, as well as a demographic questionnaire. The study has ethical approval from the Comité de protection des personnes Ouest III (number 23.03773.000242) and conformed to the Declaration of Helsinki, apart from pre-registration. Participants were paid for their time. Inclusion criteria were both right- and left-handed people above 18 years. Participants were excluded if: they had any disorder or pathology likely to affect tactile sensitivity or motor skills (e.g. epilepsy, diabetic neuropathy, dermatological condition, psychiatric or neurological disorders); were a person deprived of liberty and under legal protection, guardianship or curatorship; pregnant or breast-feeding; or experienced any type of dizziness or motion sickness, due to the AR environment.

### Experimental set-up

Participants were seated comfortably in a chair wearing an Oculus Quest 2 headset (Meta, CA), with Oculus Hand Tracking monitoring hand movements with the cameras built into the helmet (Figure 1A). The virtual environment was designed using Unity (Version 2022.3.20f1; Unity Technologies). We chose to use AR instead of VR to preserve the participant’s real-world visual context while overlaying virtual stimuli, thereby enhancing the ecological validity of the experiment. Haptic feedback was delivered via a custom 3D-printed wristband made of thermoplastic polyurethane, housing a voice-coil actuator with a resonance frequency at 133 Hz, driven by a Class-D amplifier (Bonnet et al. 2025, 2026). The wireless system, enabled precise, real-time control of vibration patterns with consistent skin contact for reliable experimental delivery. Participants wore the wristband their dominant hand, placed on the flexor tendons ~2 cm from the wrist crease (Figure 1B).

**Figure 1.**
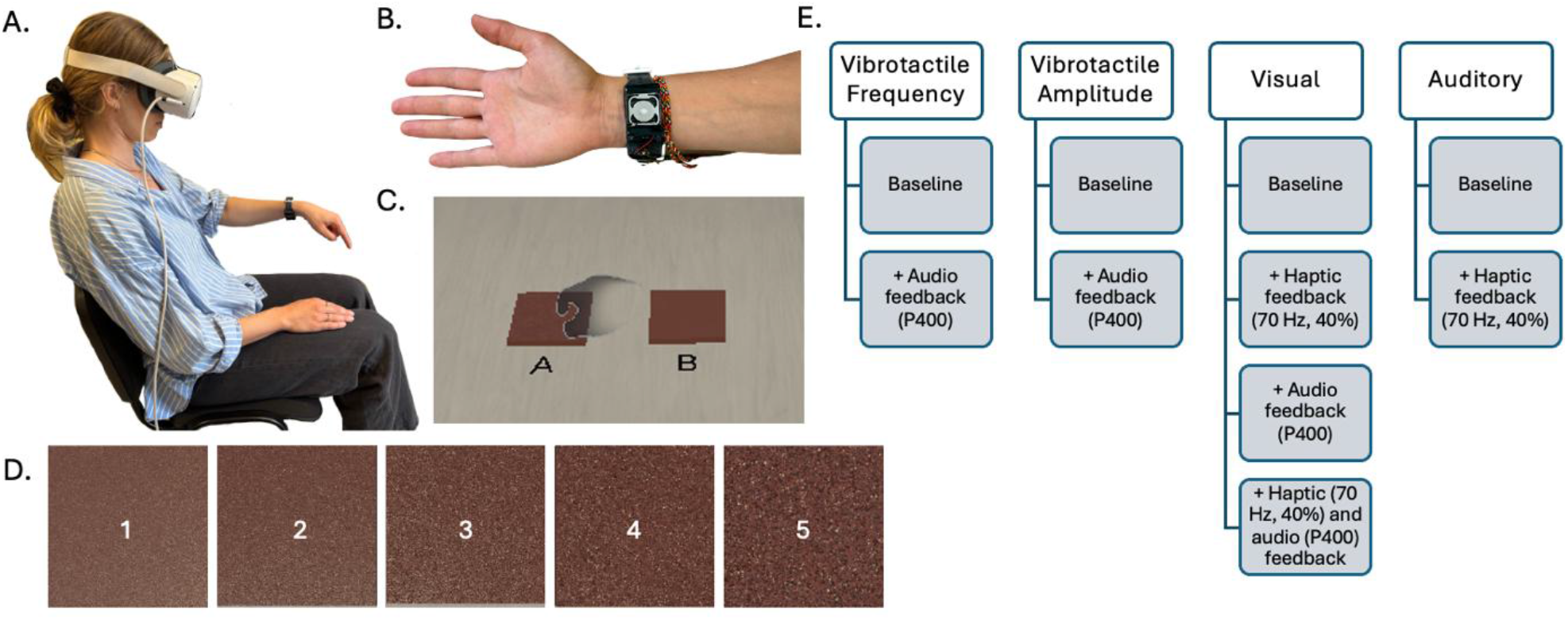
Experimental set-up. A. The participant was sat comfortably on a chair wearing the Oculus Quest 2 headset. Note that the photo is of one of the authors demonstrating the setup. B. Vibrotactile haptic feedback wristband. C. Virtual table and textures (labeled ‘A’ and ‘B’) in the experimental conditions. D. The 5 levels of visual roughness (1 = smoothest to 5 = roughest). E. Experimental design showing the four sensory dimensions and corresponding feedback conditions. In total, the experiment comprised 10 distinct conditions across the four sensory dimensions.

Participants used the index finger of their dominant hand to perform the experiment. To ensure that participants were able to readily perceive roughness differences, they first sorted five real sandpapers: P60, P240, P600, P1200, P10000 (FEPA P grading system on graining size, where a higher P corresponds to finer sandpaper), from roughest to smoothest. All participants sorted the sandpapers correctly. Before starting the experiment, participants completed a short demographic questionnaire (age, gender, handedness, previous experience of augmented and virtual reality, and haptic feedback). After the experiment, participants were asked which modality they found easiest to rely on when making the roughness judgment, and which modality that was their favorite to rely on for roughness discrimination. Next, a familiarization task was performed where participants explored six virtual textures that had different qualities in the visual, tactile, and auditory modalities. Participants were instructed to move their index finger at a constant, natural pace over the virtual texture from the top to the bottom. Textures were presented on a virtual table in front of the participants, therefore when touching the texture, their finger moved freely in the air (Figures 1A, 1C). When participants touched the virtual texture, the wristband gave a constant vibration and the roughness sound was played, via the headset, for the duration of the interaction with the texture.

### Task and stimuli

The experiment consisted of a 2-forced choice pairwise comparison between two textures labeled ‘A’ and ‘B’ where participants were instructed to stroke each texture once, starting with A that was placed on the left-hand side (Figure 1C). For each trial, the question was “Which texture is the roughest?” and responses were given verbally and noted.

To determine the range of vibrotactile parameters, we first conducted a set of pretests. Based on these results, we selected five values for each tested modality. For the frequency condition, we used 50, 60, 70, 80, and 90 Hz, all delivered at a constant amplitude of 40% (in arbitrary units given as percentages of the maximum actuator output). They were sinusoidal vibrations with respective peak-to-peak accelerations of 0.9, 1.3, 2.1, 3.0, 4.7 m/s^2^ (Bonnet et al. 2025). For the amplitude condition, we selected 40, 45, 50, 55, and 60% arbitrary units, all presented at a fixed frequency of 70 Hz. The corresponding sinusoidal peak-to-peak accelerations were 2.1, 2.4, 2.6, 2.9, 3.3, 3.4 m/s^2^, showing a linear relationship between amplitude and acceleration (Bonnet et al. 2025).

The visual stimuli consisted of sandpaper models with varying spatial frequencies, created in Blender (Version 4.3.2; Blender Online Community, 2024) to mimic different degrees of surface roughness (Figure 1C). Five textures were chosen from our pretest based on their apparent grain size. From the visually roughest to the smoothest texture, we applied scaling factors of 0.3, 0.5, 0.7, 15, and 100, categorized into five visual roughness levels from 1 = smoothest (100) to 5 = roughest (0.3). The blend file and images are available at <u>OSF</u> (https://osf.io/4r3hf). These textures were exported to the Unity game engine and standardized in size (6.5 cm^2^).

The auditory stimuli were generated by recording the sound of an index finger sliding naturally at a constant speed over five different physical sandpapers with increasing coarseness: P7000 (particle size 3 μm), P800 (22 μm), P400 (35 μm), P150 (93 μm), and P60 (260 μm), categorized into five auditory roughness levels from 1 = smoothest (P7000) to 5 = roughest (P60). This covered a large range of roughnesses from fine (particle size <20 μm) to course (particle size >100 μm) (Hollins and Risner 2000). For each sandpaper, 20 movements were recorded. The sounds were normalized, averaged, and used as stimuli delivered via the headset. The sounds are available at <u>OSF</u> (https://osf.io/4r3hf).

A total of 10 experimental conditions of different combinations of uni-modal and multi-modal stimuli were tested (Figure 1E): vibrotactile frequency, vibrotactile frequency + audio feedback, vibrotactile amplitude, vibrotactile amplitude + audio feedback, visual, visual + haptic feedback, visual + audio feedback, visual + haptic and audio feedback, auditory, auditory + haptic feedback. The conditions used the above stimuli parameters in the primary (baseline) modality and, when combined haptic or audio feedback was given in the multi-sensory conditions, haptic feedback was at 70 Hz sinusoidal vibration with an amplitude of 40%, corresponding to a peak-to-peak acceleration of 2.1 m/s^2^, while audio feedback was the P400 sound. For non-visual conditions (vibrotactile frequency, vibrotactile amplitude, and auditory), the visual aspect of the textures was always presented at roughness level 3 (medium rough).

Within each condition, participants compared all possible pairwise combinations of the five levels, resulting in 10 comparisons per condition. This yielded a total of 100 unique trials, which were repeated three times, resulting in a total of 300 trials. The order of conditions and trials was fully randomized for each participant. To reduce fatigue, participants were given two five-minute breaks after every 100 trials (approximately every 15 minutes).

### Quantification and statistical analysis

Pairwise comparison data were analyzed in Python (version 3.10) within the PyCharm IDE (JetBrains, version 2024.2.2), using the Bradley-Terry-Luce (BTL) model (Bradley and Terry 1952) to transform binary preference judgments (“A rougher than B”) into continuous latent preference scores for each stimulus. This modeling approach provides a normalized scale of relative perceptual preference, allowing direct comparison of stimuli across experimental conditions. To examine differences between parameters within conditions, we first assessed the normality of the data using Shapiro-Wilk tests. A linear regression analysis was conducted to examine the relationship between the BLT scores and the ordered parameter levels within each condition. Individual linear regressions for each participant and condition were performed to estimate the slope coefficient (β), reflecting perceptual sensitivity. The β values were then compared between conditions within each sensory dimension using Friedman tests and Wilcoxon signed-rank post-hoc tests (FDR corrected). This analysis aimed to determine whether there were differences in discrimination performance across feedback conditions. Finally, Kruskal-Wallis tests and Mann-Whitney post-hoc tests (FDR corrected) were applied to determine whether participants’ choice of ease of modalities impacted β values according to conditions.

## RESULTS

### Differences in perceived roughness between conditions

Linear regression analyses revealed strong relationships between parameter values (vibrotactile frequency, vibrotactile amplitude, visual roughness, and auditory roughness) and roughness perception across all conditions (all p < 0.01). For vibrotactile frequency conditions (Figure 2A), roughness increased linearly with frequency level, showing high model fits both in the baseline condition (slope = 9.67, r^2^ = 0.99, p < 0.001) and frequency + audio feedback (slope = 5.76, r^2^ = 0.99, p < 0.001). A similar pattern emerged for vibrotactile amplitude conditions (Figure 2B), with significant positive slopes in both amplitude baseline (slope = 1.67, r^2^ = 0.92, p = 0.010) and amplitude + audio feedback (slope = 0.62, r^2^ = 0.95, p = 0.005). For visual conditions (Figure 2C), regression analyses showed very strong linear relationships across all visual tests. Namely, in the visual baseline condition, the model fit was very high (slope = 13.40, r^2^ = 0.996, p < 0.001) and these relationships persisted with visual + audio (slope = 7.45, r^2^ = 0.998, p < 0.001), visual + haptic (slope = 6.56, r^2^ = 0.977, p = 0.0015), and visual + audio + haptic feedback (slope = 3.66, r^2^ = 0.991, p < 0.001). Finally, auditory conditions (Figure 2D) showed strong positive trends for the auditory baseline condition (slope = 3.94, r^2^ = 0.83, p = 0.031), while auditory + haptic feedback had a weaker relationship, showing a near-significant trend (slope = 1.47, r^2^ = 0.76, p = 0.054).

**Figure 2.**
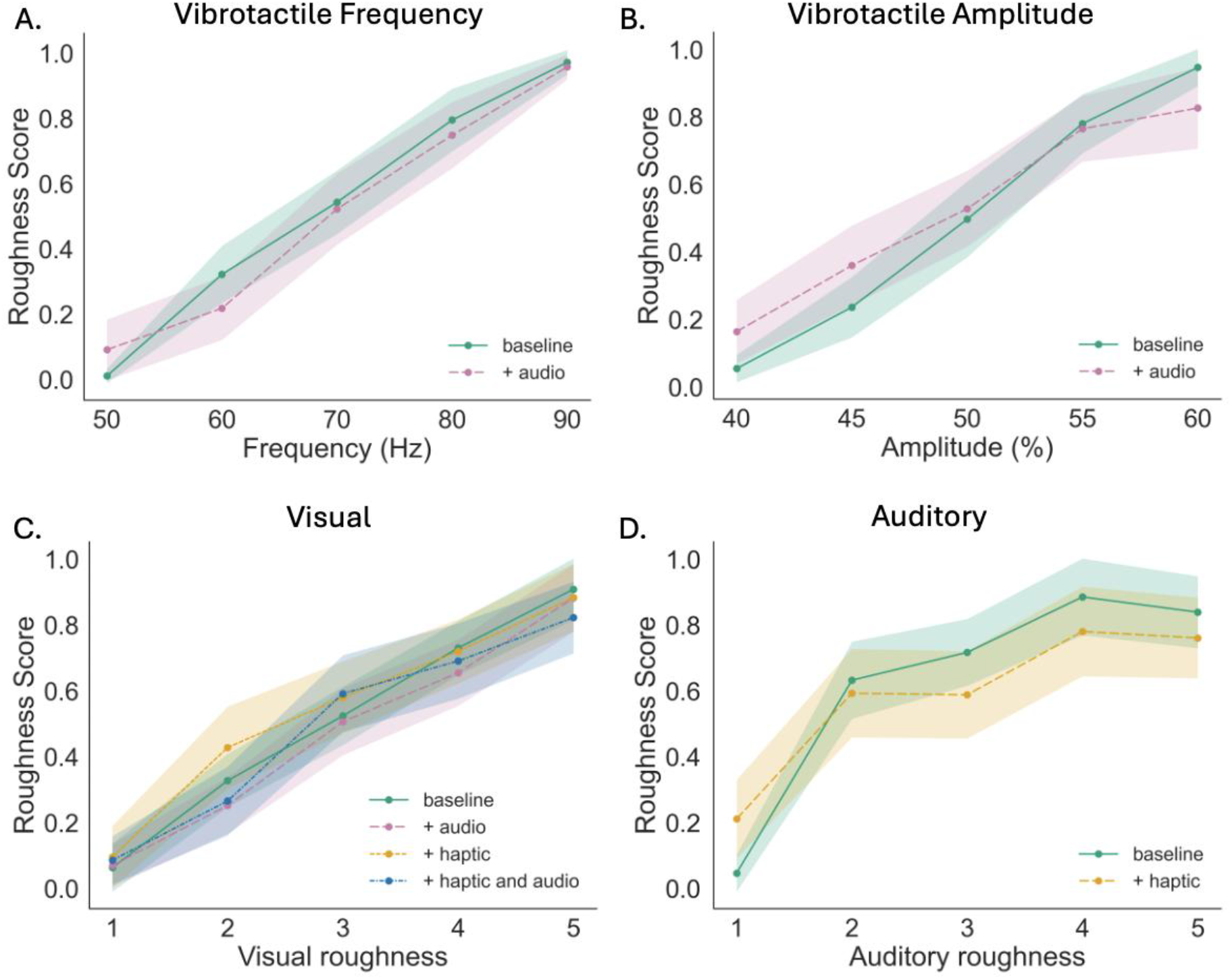
Normalized roughness scores across sensory modalities. A. Roughness scores as a function of vibration frequency (Hz), in the frequency baseline and frequency with audio feedback conditions. B. Roughness as a function of vibration arbitrary amplitude (%), in the frequency baseline and frequency with audio feedback conditions. C. Ratings across five visual roughness levels under four multisensory conditions: visual baseline, visual with haptic feedback, visual with audio feedback, and visual with haptic and audio feedback. D. Ratings across auditory roughness levels, in auditory baseline and auditory with haptic feedback conditions. Mean curves (n=30 participants) represent the average normalized BTL score across participants for each condition, with shaded bands indicating 95% confidence intervals.

### Roughness discrimination between conditions

To determine whether certain modalities were better in discriminating roughness than others, we compared the slopes of individual participant linear regressions between uni-modal baseline conditions. A Friedman ANOVA revealed a significant main effect (χ^2^(3) = 37.0, p < 0.001) indicating that the estimated beta values differed significantly among conditions, as shown in Figure 3. Subsequent pairwise Wilcoxon signed-rank tests revealed significantly lower beta values for vibrotactile amplitude compared to vibrotactile frequency (p < 0.001) and visual (p = 0.001) conditions, and lower beta values for auditory compared to vibrotactile frequency (p < 0.001) and visual (p = 0.003) conditions.

**Figure 3.**
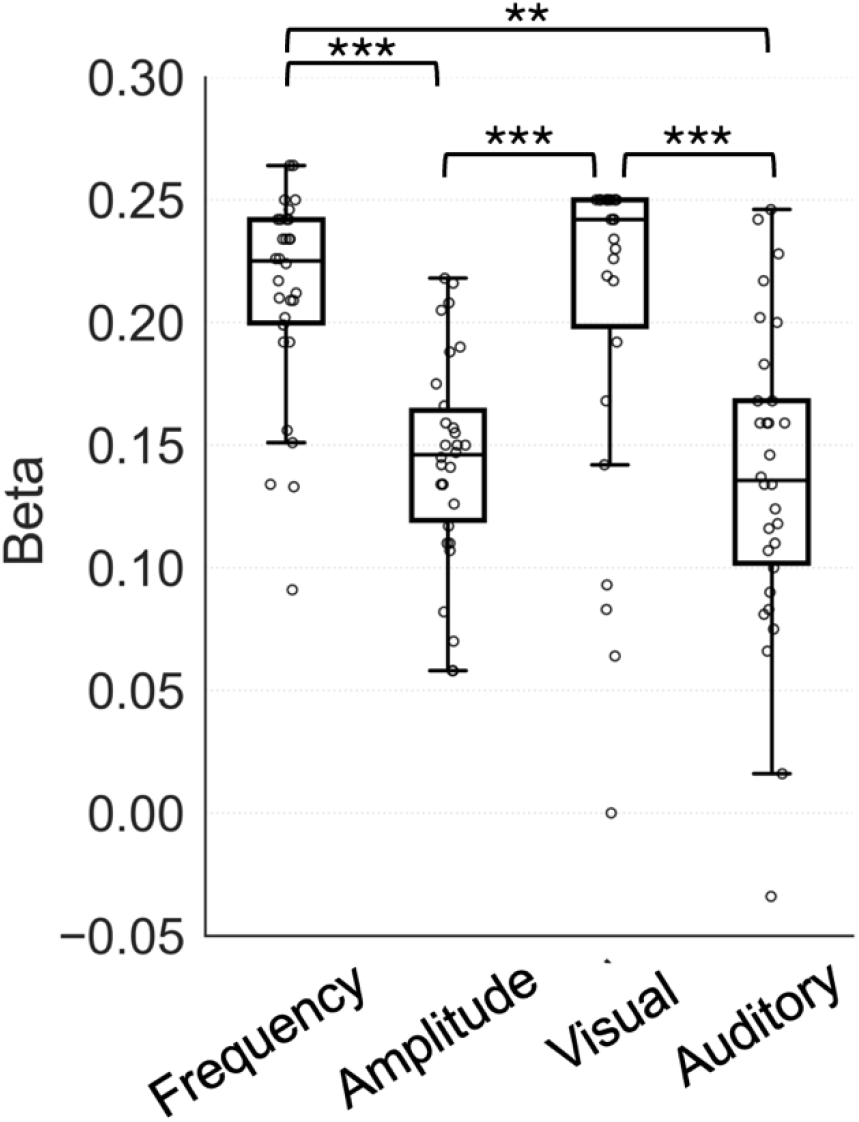
Individual beta coefficients from linear regression analyses comparing roughness sensitivity across baseline conditions. The box plot shows the distribution of beta values across the 30 participants for the baseline conditions (without any extra feedback). Amplitude and auditory show statistically lower beta values than frequency and visual, revealing weaker roughness discrimination. The dots represent individual data points. Significance levels: **p < 0.01, ***p < 0.001.

Using the same individual linear regression slopes (β), we analyzed whether perceptual relationships varied significantly between all conditions per modality. For vibrotactile frequency (Figure 4A), a Wilcoxon test indicated a significant decrease in perceptual sensitivity from the frequency baseline to the frequency + audio feedback condition (W = 61.0, p = 0.001). For vibrotactile amplitude (Figure 4B), a significant reduction was also found from the amplitude baseline to the amplitude + audio feedback condition (W = 109.5, p = 0.020). For the visual dimension (Figure 4C), the Friedman test yielded a significant main effect (χ^2^(2) = 24.7, p < 0.001), demonstrating that perceptual sensitivity in visual conditions differed significantly between tested configurations. Post-hoc Wilcoxon comparisons showed significant differences in slope values for all individual comparisons between conditions (p < 0.05), apart from one comparison, namely, visual + haptic feedback and visual + haptic and audio feedback comparisons. This result suggests reduced perceptual slopes when haptic and/or audio feedback was added to the visual condition. For the auditory condition (Figure 4D), a significant decrease in slope values was found from the auditory baseline, as compared to the auditory + haptic feedback (W = 39.0, p < 0.001).

**Figure 4.**
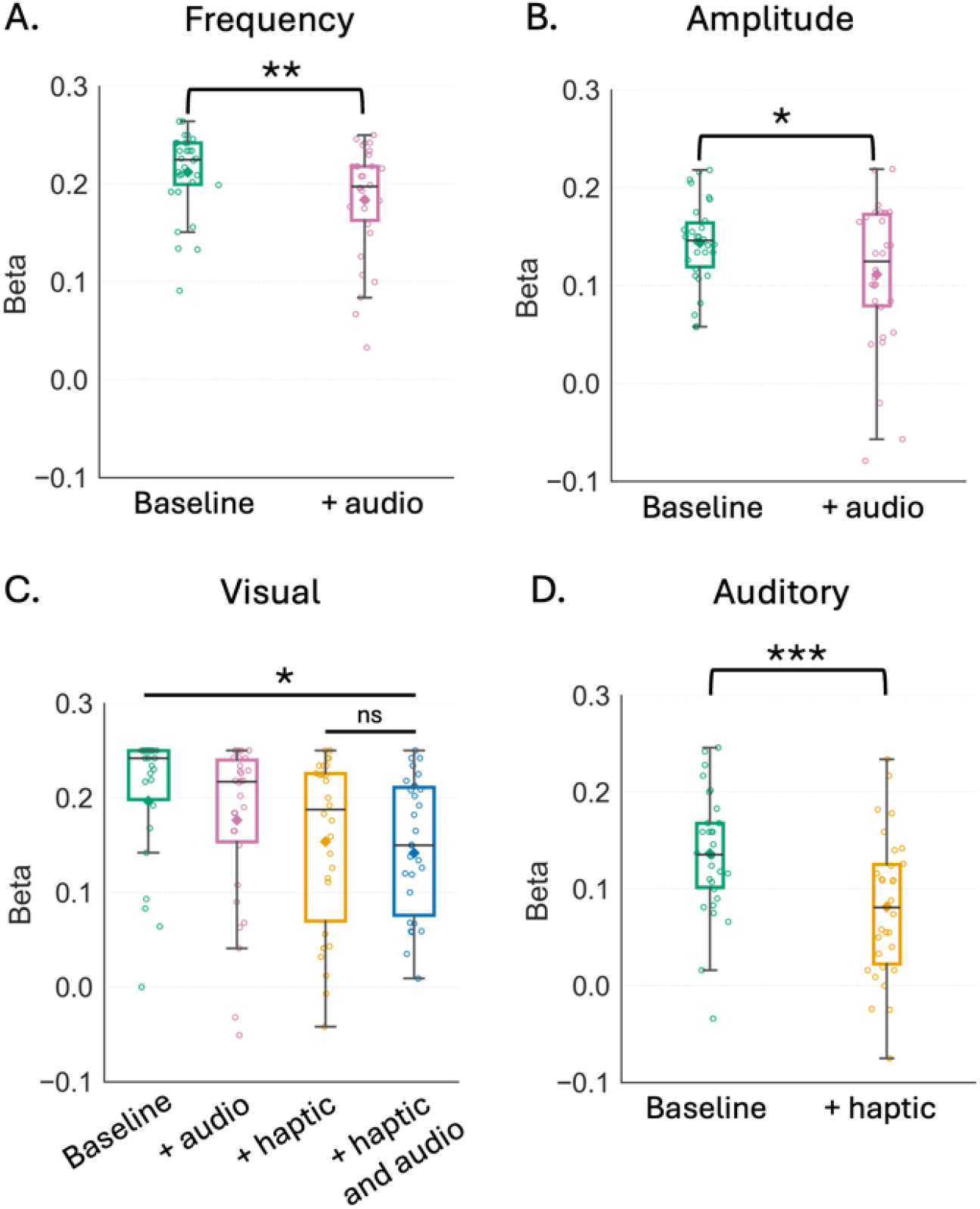
Individual beta coefficients from linear regression analyses comparing roughness sensitivity across sensory feedback conditions. Each box plot shows the distribution of beta values across the 30 participants for the following conditions: A. frequency baseline and frequency + audio feedback, B. amplitude baseline and amplitude + audio feedback, C. visual baseline, visual + haptic feedback, visual + audio feedback, visual + haptic and audio feedback, where all comparisons were significant apart from the one comparison marked ns, and D. auditory baseline and auditory + haptic feedback conditions. Significance levels: ns - not significant, *p < 0.05, **p < 0.01, ***p < 0.001.

### Participant preferences for discriminating roughness

The preferences questionnaire revealed that in order to discriminate roughness, 40% of participants found the haptic dimension easier to rely on, 37% found the visual dimension easier, while 23% found the auditory dimension easier. To examine whether participants’ perceptual trends varied depending on which modality they perceived as easiest to rely on when making the roughness comparison, we analyzed linear regression slopes (β) across easier modalities for each condition. Significant effects of perceived ease were found in several conditions (Figure 5). In the vibrotactile frequency and the vibrotactile frequency + audio feedback conditions, significant main effects were observed (H(2) = 7.63, p = 0.022, and H(2) = 10.41, p = 0.005, respectively). Post-hoc comparisons revealed higher beta values (i.e. perceptual sensitivity) in participants who reported the haptic modality as easiest than those who found the visual modality easiest (p = 0.02 for vibrotactile frequency condition and p = 0.015 for vibrotactile frequency + audio feedback condition).

**Figure 5.**
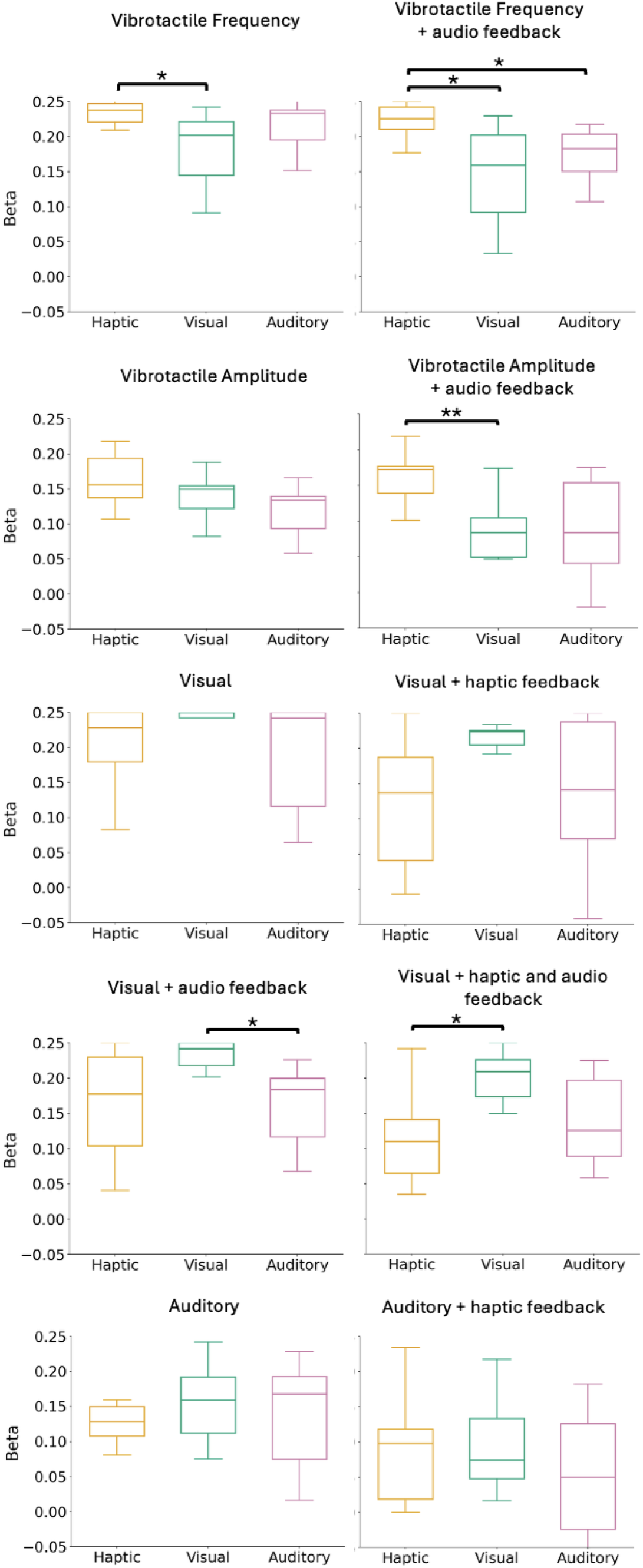
Individual beta coefficients from linear regression analyses grouped by the sensory modality participants reported as the easiest to rely on (Haptic, Visual, Auditory). Each box plot shows the distribution of beta values among 30 participants for all conditions. Only the vibrotactile frequency baseline, vibrotactile frequency + audio feedback, vibrotactile amplitude + audio feedback, visual + audio feedback, and visual + audio and haptic feedback conditions show significant differences between modalities. Each box represents the distribution of individual beta values among participants within each modality. Higher beta values reflect greater sensitivity to stimulus variation. Significance levels: *p < 0.05, **p < 0.01, ***p < 0.001.

These results suggest that perceived ease modulated sensitivity in frequency-based roughness judgments. The same pattern was found in the vibrotactile amplitude with audio feedback condition, where a significant effect of ease of perception was found (H(2) = 12.08, p = 0.002), reveling significantly higher beta values for participants who ranked haptic as the easiest modality compared to visual (p = 0.002). Moreover, a significant effect was observed in the visual + audio feedback condition (H(2) = 6.87, p = 0.032), showing higher beta values for participants who preferred visual versus auditory modalities (p = 0.041). The visual + haptic and audio feedback condition was also significant (χ^2^(2) = 6.65, p = 0.036), where participants who chose the visual modality had higher beta valuer than those who preferred the haptic modality (p = 0.045).

## DISCUSSION

The present study examined how roughness perception can be modulated through vibrotactile, visual, and auditory cues in an augmented reality environment. Perceived roughness showed strong linear relationships with stimulus parameters across all modalities, with vibration frequency and visual granularity yielding the highest discrimination performance, while auditory cues were less consistent. Crucially, the addition of non-informative sensory feedback systematically degraded roughness discrimination, suggesting that task-irrelevant inputs introduce perceptual noise in AR. Individual sensory preferences further modulated these effects, with haptic-reliant participants showing greater sensitivity to vibrotactile variations and visually-reliant participants demonstrating stronger responses to visual cues. Together, these findings highlight the multimodal nature of roughness perception in virtual environments, emphasize the importance of individual differences in multisensory integration (Proulx et al. 2022; Ward 2019), and offer concrete insights for implementing adaptive, immersive multisensory systems.

In our haptic vibrotactile conditions, perceived roughness increased linearly with both vibration frequency and amplitude. Classical touch theories distinguish an intensive coding based on vibration amplitude, and a temporal coding based on frequency, predicting that roughness increases with amplitude but decreases as frequency rises (Bensmaia and Hollins 2003). In our study, however, roughness increased with frequency across the tested range (50-90 Hz). This finding is consistent with our previous work showing that perceived vibration intensity at the wrist increases with both frequency and amplitude and is strongly correlated with vibratory acceleration (Bonnet et al. 2025). The apparent contradiction with classical models is likely explained by the specific frequency range employed, which falls precisely in the transition zone between FA1 (Meissner) and FA2 (Pacinian) afferent sensitivity (Talbot et al. 1968), which is between fine and coarse textures. In this intermediate range, increases in frequency coincide with higher perceived vibratory intensity, driven by the sharp rise in Pacinian afferent sensitivity above approximately 60 Hz. A change in frequency in this band therefore does not function as a purely temporal cue but is physiologically equivalent to a change in intensity, consistent with Pacinian-weighted intensity models (Bensmaïa and Hollins 2005). The strong implication of FA2 afferents is further supported by the markedly higher discrimination performance observed for the frequency condition compared to amplitude or auditory conditions. Unlike amplitude, which is the dominant factor in fine-texture roughness at the fingertip (Bensmaia and Hollins 2003), vibrotactile stimulation applied remotely at the wrist radically alters sensory dynamics: FA2 afferents, which are sensitive to remote vibrations, likely drive discrimination performance when spatial cues from the fingertip are absent (Romo et al. 2002; Talbot et al. 1968). We therefore postulate, frequency variations were relied upon to infer intensity, preferentially transmitted via the Pacinian channel.

Visual roughness perception similarly showed strong positive linear relationships with stimulus parameters, and visual discrimination performance was equivalent to that observed for vibrotactile frequency, as well as being significantly superior than vibrotactile amplitude and auditory conditions. These findings extend prior work on visual texture cues (Ho et al., 2006; Klatzky & Lederman, 2010) and confirm that roughness can be systematically controlled through visual grain size, even in AR, without any tactile input.. In line with our results, it has been found that vision and touch perception give similar visuotactile properties, even showing similar performances in roughness judgements (Binns 1937; Björkman 1967; Lederman and Abbott 1981).

Regarding auditory conditions, listening to friction sounds from sandpaper of varying granularity produced a linear increase in roughness perception, though with a lower discrimination slope than vibrotactile frequency or visual conditions. Auditory cues can modulate the perception of roughness (Guest et al. 2002; Suzuki et al. 2008) and our results align with the finding that auditory cues are given less weight in a multisensory context (Di Stefano & Spence, 2022; Klatzky & Lederman, 2010). Furthermore, as audition is not the dominant modality for discriminating roughness arising in tactile interactions, this was reflected in our finding that only 23% of participants found the auditory presentation the easiest to rely on for roughness discrimination.

Participants who rated a specific modality as easiest to rely on for roughness discrimination also demonstrated increased perceptual sensitivity within that modality. This pattern is consistent with the modality appropriateness hypothesis, whereby the modality best suited for a given individual in a task tends to dominate perception (Weisenberger and Poling 2004). In the context of texture perception specifically, haptic and visual modalities are generally considered the most appropriate (Baumgartner et al. 2013; Lederman 1979), which aligns with our finding that participants relying on these two modalities showed the clearest sensitivity advantages. This is further consistent with evidence that individuals construct stable but idiosyncratic internal representations of roughness across modalities (Bergmann Tiest and Kappers 2007). The existence of at least two perceptual subgroups, haptic-reliant and visually-reliant users, has direct implications for AR system design. Rather than applying uniform multisensory feedback, future systems should implement sensory profiling to dynamically tailor feedback weighting to the individual user’s perceptual strategy, thereby maximizing both discrimination accuracy and subjective realism.

A consistent finding across all modalities was that adding extra sensory feedback that carried no discriminative information between the two textures systematically reduced perceptual sensitivity. This pattern suggests that task-irrelevant inputs function as perceptual noise, interfering with roughness discrimination. This is supported by the weighted averaging model of multisensory integration, whereby even non-informative cues can disrupt dominant sensory signals (Parise and Ernst 2017). The effect may also reflect a binding problem: temporally synchronous, but semantically incongruent, inputs may be automatically integrated into a unified percept, introducing perceptual ambiguity (Stein and Stanford 2008). Furthermore, the cognitive resources required to resolve such incongruence likely imposed additional processing demands, impairing performance in line with cognitive load theory (Sweller 1988). For AR design, this finding has direct practical implications, where adding non-informative multisensory feedback would actively degrade the perceptual experience. Multisensory feedback should therefore be carefully curated to ensure that every added cue carries task-relevant, congruent information.

Although promising for AR applications, the present study has limitations. In everyday interactions, sounds generated by skin-surface contact are more subtle and non-salient compared to other sensory inputs, with vision and touch typically dominating surface property judgments (Guest and Spence 2003b, 2003a; Lederman 1979). However, our auditory feedback did not come directly from the virtual touch, but occurred via the headset, meaning a potential mismatch in sound location and realism. Although the phenomenon of spatial ventriloquism can facilitate the perceptual localization of sounds to co-occurring visual events (Chen & Vroomen, 2013), this spatial discrepancy between the expected and actual sound source may have weakened multisensory binding coherence, which may partly account for the relatively weaker and more variable effects observed in the auditory conditions. Second, the exact playback dynamics of the auditory stimuli were not fully matched to small variations in finger movement, rather the sound was played on movement. Such small temporal inconsistencies could disrupt the fine-grained alignment between tactile and auditory signals that is known to be critical for effective multisensory integration (Guest et al. 2002). Future implementations could use adaptive sound synthesis driven by real-time movement parameters to preserve temporal congruence across modalities. Third, our sample consisted predominantly of young adults, which may limit the generalizability of the findings to other populations. Age-related changes in tactile sensitivity, sensory integration efficiency, and processing speed (Samain-Aupic et al. 2024; Stevens 1992) are well-documented and could alter how multisensory feedback is weighted and perceived in virtual environments, meaning parameters should be adapted to individuals.

In conclusion, the present work advances our understanding of multisensory roughness perception in AR by demonstrating that haptic, visual, and auditory cues can each independently and systematically modulate perceived roughness, while also revealing that non-informative feedback degrades perceptual performance. These findings have direct implications for the implementation of immersive systems: multisensory feedback should be carefully curated for congruence and task-relevance, grounded in individual sensory profiling. Future research should expand the range of stimulus parameters and examine how users resolve conflicts between incongruent haptic, visual, and auditory cues. Longitudinal designs could address whether individual sensory strategies are stable across tasks or can be shaped through training and exposure. Finally, extending this paradigm to broader populations, including older adults and individuals with sensory impairments, is essential to ensure that the multisensory design principles identified here translate into inclusive and generalizable AR systems. Together, these directions point toward a future in which immersive environments are not only visually convincing, but perceptually coherent across all the senses.

## RESOURCE AVAILABILITY

### Lead contact

Requests for further information and resources should be directed to and will be fulfilled by the lead contact, Rochelle Ackerley.

### Data and code availability

- Data reported in this paper has been deposited at OSF at https://osf.io/4r3hf and is publicly available as of the date of publication.
- All original code reported in this paper will be shared by the lead contact upon request.
- Any additional information required to reanalyze the data reported in this paper is available from the lead contact upon request.

## ACKNOWLEDGMENTS

This work funded by an industrial CIFRE PhD thesis bursary to S. Bonnet from Haptify in collaboration with the Centre National de la Recherche (CNRS).

## AUTHOR CONTRIBUTIONS

Conceptualization, S.B., H.D., and R.A.; methodology, S.B. and R.A.; Investigation, S.B., R.E.; writing – original draft, S.B.; writing – review & editing, S.B. and R.A.; funding acquisition, H.D. and R.A.; resources, S.B., I.Z. and H.D.; supervision, H.D. and R.A.

## DECLARATION OF GENERATIVE AI AND AI-ASSISTED TECHNOLOGIES IN THE WRITING PROCESS

During the preparation of this work, the author(s) used Emmy, developed by Mistral AI for the CNRS, in order to improve the fluency of the English language during the drafting stage of the paper. After using this tool or service, the author(s) reviewed and edited the content as needed and take(s) full responsibility for the content of the publication.

## KEY RESOURCES TABLE

| REAGENT OR RESOURCE | SOURCE | IDENTIFIER |
| --- | --- | --- |
| Software and algorithms |  |  |
| Python version 3.10 | Python Software Foundation | https://www.python.org/ |
| Unity 2022.3.20f1 | Unity Technology | https://unity.com/ |
| Meta Quest 2 SDK | Meta Platforms | https://developer.meta.com/ |
| Python version 3.10 | Python Software Foundation | https://www.python.org/ |
| PyCharm 2024.2.2 | JetBrains | https://www.jetbrains.com/pycharm/ |
| Blender 4.3.2 | Blender Online Community | https://www.blender.org/ |
| Other |  |  |
| Supplementary materials | OSF | [OSF](https://osf.io/4r3hf) (https://osf.io/4r3hf) |

